# Direct calculation of cryo EM and crystallographic model maps for real-space refinement

**DOI:** 10.1101/2022.07.17.500345

**Authors:** Alexandre Urzhumtsev, Ludmila Urzhumtseva, Vladimir Y. Lunin

**Affiliations:** Centre for Integrative Biology, Institut de Génétique et de Biologie Moléculaire et Cellulaire, CNRS–INSERM-UdS, 1 rue Laurent Fries, BP 10142, 67404 Illkirch, France; Université de Lorraine, Faculté des Sciences et Technologies, BP 239, 54506 Vandoeuvre-les-Nancy, France; Architecture et Réactivité de l’ARN, UPR 9002 CNRS, IBMC (Institute of Molecular and Cellular Biology), 15 rue R. Descartes, 67084 Strasbourg, France; Université de Strasbourg, Strasbourg, F-67000 France; Institute of Mathematical Problems of Biology RAS, Keldysh Institute of Applied Mathematics of Russian Academy of Sciences; Pushchino, 142290, Russia

**Keywords:** Real-space refinement, atomic images, resolution, atomic displacement parameter, Fourier ripples, truncation distance, interference function, shell decomposition

## Abstract

Real-space refinement of atomic models improves them by their fit to experimental scattering electrostatic potential maps in cryo electron microscopy and to electron density maps in crystallography. This procedure has a number of advantages in comparison with reciprocal-space refinement, is complementary to it in crystallographic studies and is the principal technique in cryo EM. An accurate real-space refinement of atomic models can be done by comparison of the model maps, calculated according to the respective theory, with the experimental ones, when the former mimic imperfections of the latter, mainly a limited resolution and an atomic disorder. Calculation of model maps as a sum of contributions of individual atoms means that these contributions - atomic images in the given map – should also be affected by the resolution and positional disorder. This blurs atomic images and surrounds their central peak by Fourier ripples. These ripples can be described by a specially designed function which allows combining both principal effects of map imperfection in an analytic way. The atomic images, at any resolution and with any value of the atomic displacement parameter, can be decomposed into a linear combination of such functions with the precalculated parameter values. As a consequence, each map value becomes an analytic function of atomic parameters including displacement parameter and a local resolution if required. Using the chain rule for the score function which compares the maps, such model results in analytic expressions for its partial derivatives with respect to all atomic parameters allowing an efficient real-space refinement. At the same time, for practical calculations atomic images are cut at some truncation distance, *i.e*., include a limited number of ripples. This introduced in the model maps errors which are not present in the experimental maps. Due to oscillating behavior of the atomic images, the choice of the value of this truncation distance is not straightforward and discussed in this work.

**Synopsis:** A new method is suggested to calculate maps of a limited and eventually inhomogeneous resolution as an analytic function of all atomic parameters, including local resolution.

## 1. Introduction

In Introduction, we remind several known results. We do this due to existing confusing between qualitatively similar entities while the difference between them makes major issues in the practical problems discussed below.

### 1.1. Real-space refinement and density maps

Quality of a macromolecular atomic model defines how meaningful the model is. Initial models are constructed in the maps, in most cases obtained by X-ray or neutron diffraction experiment or by cryo-electron microscopy (cryo-EM). Even when recently power tools have been suggested to predict spatial protein structures (Jumper *et al*., 2021; Baek *et al*., 2021), refinement of atomic models by their comparison with the respective experimental maps is the crucial step to get reliable and accurate information about the structure of macromolecules and their complexes (Palmer & Aylett, 2022; Helliwell, 2022).

These maps represent different scalar fields: electron density distribution, nuclear density, electrostatic potential or electrostatic scattering potential. To make it short, in what follows we call all of them as density distributions, *ρ_experim_*(**r**). The experimental maps are images of the respective fields. As a rule, they do not show an atomic structure as ‘point’ atoms and suffer from imperfections that are mainly caused by a positional disorder of the structure under study, both dynamic and static, and by the limited resolution of the maps. Eventually, their resolution may be different in different map regions leading to the concept of the local resolution (Kucukerlbir *et al*., 2014; Vilas *et al*., 2018; Ramírez-Aportela *et al*., 2019); this is especially true for the maps obtained by cryo EM (Cardone *et al*., 2013). Real-space macromolecular refinement, originated by Diamond (1971), improves the values of the available atomic model parameters by reducing the discrepancy between two maps, the experimental one and that calculated from the current model according to the accepted theory. To compare these two maps quantitatively, the latter map should mimic imperfections of the former. Alternatively, one may refine a point-atoms model (Afonine, Poon *et al*., 2018), which does not need a model map. However, it is applicable rather at early and intermediate stages and cannot deal with atomic displacement parameters.

We remind that map comparison used for model refinement is different from a usual visual map analysis and that their straightforward comparison can be misleading (Urzhumtsev *et al*., 2014). First, for a visual analysis one rather looks the contours shown with some cut-off level while refinement compares the map values. Second, the fit of maps means not only fit of their positive values but also negative values which should be present in the exact maps of a limited resolution (Lunin, 1988) and which do contain structural information (Urzhumtsev *et al*., 1989). Third, often maps are compared not only close to the atomic centers but in larger regions. This results in comparison of many near-zero values in the solvent region which bias the target values. Concluding, for an appropriate model refinement, the model map should be calculated more accurately than the visual analysis implies.

Atomic composition of macromolecules means that the field *ρ_experim_*(**r**) can be considered as a sum of densities from the atoms which compose the sample. Similarly, the maps of these densities can be represented by a sum of respective atomic contributions (see, for example, Diamond, 1971, Lunin & Urzhumtsev, 1984; Chapman, 1995; Mooij *et al*., 2006; Chapman *et al*., 2013; DiMaio *et al*., 2015; Sorzano *et al*., 2015; and references therein). In order to get such maps sufficiently accurate, several theoretical and practical questions should be answered. We address them in this work. The key tool of the solution is a function specially designed for these goals (Urzhumtsev & Lunin, 2022; Urzhumtseva *et al*., 2022).

### 1.2. Real and virtual crystals

Macromolecular structural studies deal with two principal types of objects. First, methods like SAXS and Cryo Electron Microscopy (cryoEM) consider objects isolated in space (single particle). Second, macromolecular crystallography (MX) considers objects in a crystal form. The latter are thought as an infinite set of the identical objects placed periodically in space. We leave aside more exotic situation such as two-dimensional crystals or nanocrystals where a too small size of crystals starts to play a role. In the principal structural methods (except, *e.g*., NMR), experiments deal with different kinds of scalar fields and with respective diffraction data. These two kinds of data are related to each other by the Fourier transform. By computational convenience, a single particle is often considered belonging to a virtual crystal with a large unit cell which contains only one isolated particle not interacting with its neighbors. In what follows we do not distinguish between real and virtual crystals.

A three-dimensional crystal is defined by three basis vectors {**a, b, c**}. They define the infinite real-space lattice

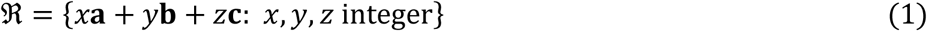

and the crystal unit cell

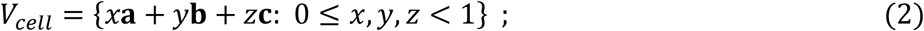

with its volume |*V_cell_*|. A scalar field *ρ*(**r**) defined in this crystal satisfies the periodicity condition

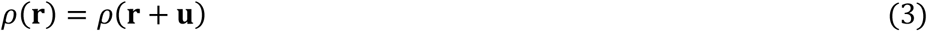

for any 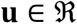 and any **r**.

### 1.3. Atomicity and direct maps

A density distribution in a crystal is considered as a sum of density distributions of individual atoms in positions **r**_*n*_

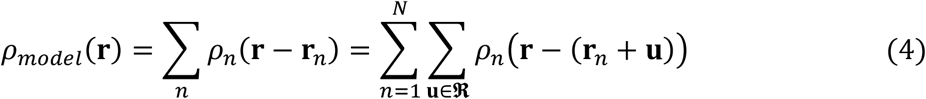

The sum is taken over the whole infinite set of the atoms in the crystal where an independent subset is composed of *N* atoms. Atomic densities are peaky functions which value decreases with the distance and become negligibly small beyond some limit, of order of a few Ångstroms. By this reason, knowing an independent set of atoms, one can calculate an approximation *ρ_direct_* (**r**) to *ρ_model_* (**r**) in any point of the crystal by a direct summation (4) over a finite subset of atoms ignoring those at a distance beyond some truncation limit from the considered point **r**. In (4) we ignore the contribution of non-structured crystal components including bulk solvent (Rupp, 2013; Urzhumtsev & Lunin, 2019; Afonine *et al*., 2021).

### 1.4. Reciprocal space representation

There exists an alternative way to calculate *ρ_model_*(**r**) as a finite number of atomic contributions. First, we define a basis {**a*, b*, c\***} dual to the basis vectors {**a, b, c**} and called the reciprocal-space basis. Like the real-space basis, the reciprocal-space basis {**a*, b*, c\***} also defines a grid:

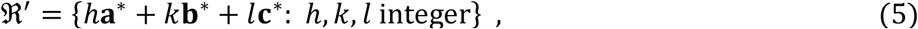

called the reciprocal space lattice.

A periodic function *ρ_model_* (**r**) may be presented by the infinite Fourier series

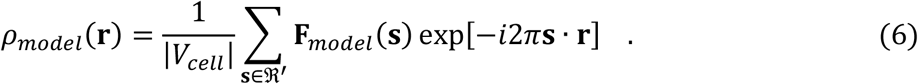

Here **s · r** is the dot (scalar) product of two vectors, and the complex-valued Fourier coefficients **F**_*model*_(**s**), called also structure factors, are calculated as the integral over the unit cell

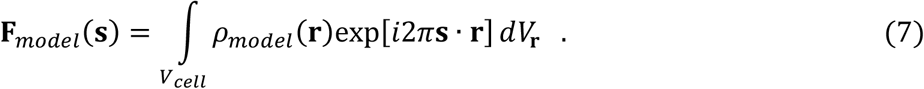

Since the Fourier transform is a linear operation, and *ρ_model_*(**r**) is a sum of atomic contributions, its structure factors **F**_*model*_(**s**) can be also calculated as a sum but this time only over an independent set of atoms, *e.g*., those inside the unit cell

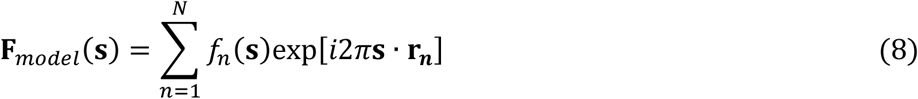

Here, a scattering function *f_n_*(**s**) of the atom *n* is calculated as the integral Fourier transform of the atomic density *ρ_n_*(**r**) over the whole space

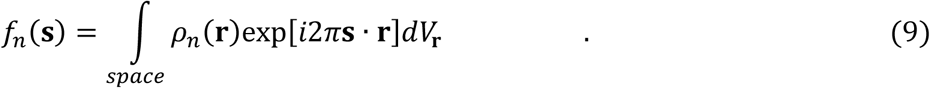

When *ρ_n_*(**r**) is spherically symmetric, its scattering function *f_n_*(**s**) is real-valued and spherically symmetric. For such functions, we note their radial components by an overline, *e.g*., 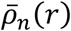 and 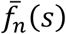 with *r* = |**r**| and *s* = |**s**|, so that 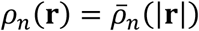. The density of an isolated atom is expressed through its scattering function as

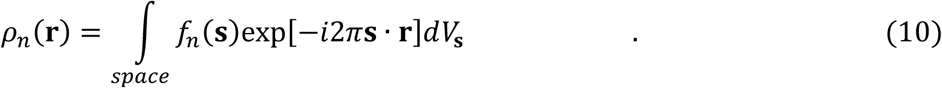

In what follows, we call a model map *direct* and note it *ρ_direct_*(**r**) if it is calculated explicitly as a sum (4) of atomic contributions in real space and call it a *Fourier map* and note it *ρ_Fourier_*(**r**) if it is calculated as the Fourier series (6) using the model structure factors.

### 1.5. Resolution cut-off and its effects

The term ‘resolution’ is defined differently in different branches of science, and even in the same branch such as in crystallography (Urzhumtseva *et al*., 2013) or cryo EM (Afonine, Klaholz *et al*., 2018). The concept mostly used in biological crystallography is the following. Being a function of a space vector **r**, a Fourier harmonic exp[-*i*2*π***s** · **r**] can be considered as a kind of a standing plane wave in the direction s with the period *d* = |**s**|^−1^. The value *d* is referred to as the resolution of the reflection corresponding to the scattering vector **s**. A Fourier synthesis 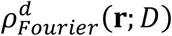 is calculated at the resolution *D* if the respective sum includes all, or ‘almost all’ (Urzhumtseva *et al*., 2013), nodes **s** of the reciprocal space lattice with |**s**| ≤ *D*^−1^, *i.e*., all reflections of the individual resolution *d* ≥ *D*:

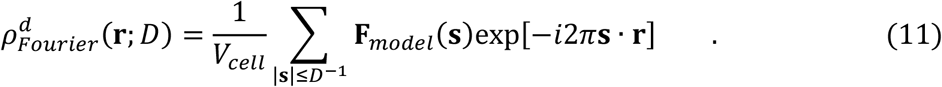

The respective direct map can be calculated as a sum

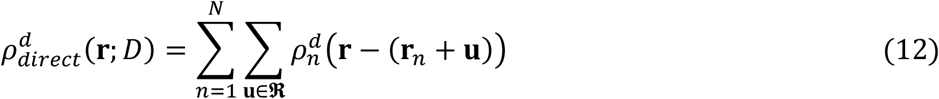

of atomic images of the same resolution *D*:

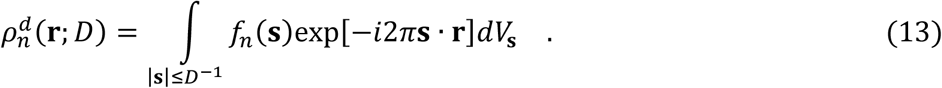

We call (13) a ‘*D* -resolution image of atom *n*’. For the simplest case of a ‘point’ atom, density of which is represented by a three-dimensional Dirak function *δ*(**r**), its scattering function is equal to one everywhere and the scattering function of its *D*-resolution image is the same but cut out beyond the sphere of the radius *D*^−1^. The image of such virtual point atom can be expressed analytically

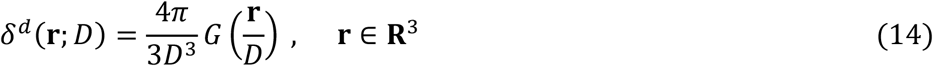

through the interference function of a dimensionless parameter **x** = *D*^−1^**r**:

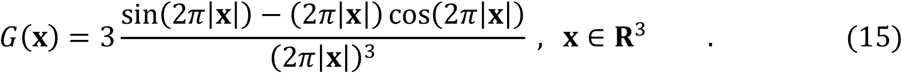

This is a spherically symmetric oscillating function of the distance *x* = |**x**| with the amplitude decreasing as *x*^−2^. Integration (13) over a sphere |**s**| ≤ *D*^−1^ can be also seen as an integration over the whole space of the same function multiplied by the three-dimensional step function equal to zero beyond |**s**| ≤ *D*^−1^. Referring to the convolution theorem, an image of any atom at a resolution *D* can be expressed as the convolution

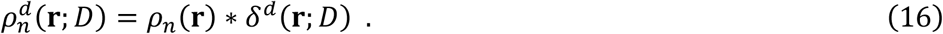

and may be assumed as the delta-function image filtered with window of the function *ρ_n_*(**r**) or *vice versa*.

### 1.6. Harmonic disorder and atomic images

Another source of imperfection of density maps is a positional disorder. Atomic positions vary about their mean values **r**_*n*_ in time and in space, from one unit cell to another in real crystals, from one image to another when working with cryo-EM projections. Except large-scale alternative conformations, such disorder is considered as a harmonic, isotropic for most of macromolecular studies. This means that its influence on the density distribution can be modeled by a convolution

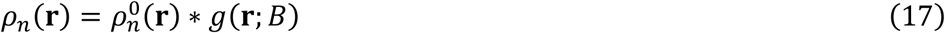

of the atomic density 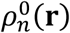 of an immobile atom with a three-dimensional isotropic Gaussian function

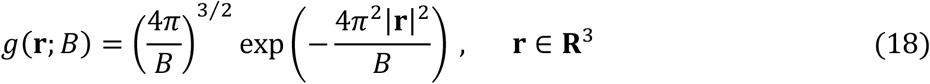

Parameter *B* is known as the isotropic atomic displacement parameter.

In reciprocal space, convolution (17) is equivalent to the product of the respective scattering function 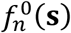 of an immobile atom by a Gaussian:

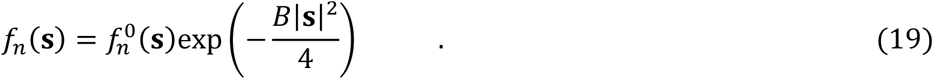

In overall, with (16), the *D*-resolution image of an atom *n* with a displacement factor *B* becomes

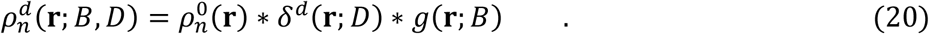

In particular, according to the properties of the Gaussian functions, an image of a ‘Gaussian atom’ 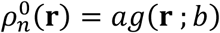 is

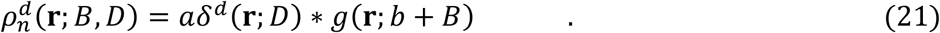

Due to the peaky shape of atomic densities, it is common to approximate them, or rather respective atomic scattering functions, by a linear combination of origin-centered Gaussians (*e.g*., Doyle & Turner, 1968; Agarwal, 1978; Waasmaier & Kiefel, 1995; Peng, 1999; Grosse-Kunstleve *et al*., 2004; Brown *et al*., 2006). This makes a virtual Gaussian atom a convenient tool for further analysis. In such case, the key problem in fast calculation of the atomic image becomes the analytic calculation of (21).

### 1.7. Map calculation method and real-space refinement

Calculated with the exact scattering functions and with no rounding errors, 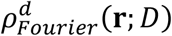; is equal to 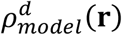 of the resolution *D*. While in crystallography it is supposed that the map resolution is the same in all its points, this is not the case for the cryo EM maps where the resolution *D* may strongly vary from one region to another (Cardone *et al*., 2013) making the concept 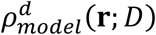 inappropriate. Another difficulty is that calculation of the model structure factors as (8) is very time consuming and they are calculated usually by the two-step procedure generating first the density distribution in a fine grid and calculating then its discrete Fourier transform (Sayre, 1951; Ten Eyck, 1977; Navaza, 2002; Afonine & Urzhumtsev, 2004) which introduces extra errors. To avoid summation over infinite number of atoms at the first step, the atomic densities are truncated at some reasonable distance *r_max_*. The total procedure requires two Fourier transforms for a map of a given uniform resolution. Finally, errors in a part of a model are propagated over all structure factors and thus over the whole cell.

Calculation of 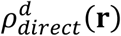 does not have these disadvantages, and its resolution can be adapted from point to point, varying from atom to atom (Chapman *et al*., 2013; Urzhumtsev & Lunin, 2022). Theoretically, 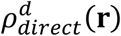 coincides with 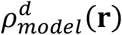 when calculated with the whole infinite sets of atoms and an infinite truncation distance of atomic images which is impossible in practice. The truncation of composing atomic images is the principal source of errors in 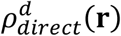, but increasing the truncation distance increases the CPU time. Another computational difficulty of the direct map calculation is an absence of analytic expressions for the atomic images (21) which makes it difficult to vary their parameters during refinement.

Below we discuss the problems of the direct map calculation and their solution. We compare different metrics and their efficiency to answer these questions. We show first that the truncation radius should be larger than often expected and requires including several Fourier ripples into calculation. Then we suggest a new method to obtain atomic images expressed by analytic functions of the coordinates and the displacement parameters. Finally, we discuss how to make such images depending analytically on the resolution as well. In overall, we suggest the procedure, including the parameters choice, to generate density maps of any resolution, even inhomogeneous over the cell. Analytical dependence of the map values of atomic parameters opens a way to a ‘real’ real-space refinement in macromolecular crystallography and in cryo EM.

## 2. Atomic images and Fourier ripples

### 2.1. Overall considerations

While, like atomic densities 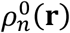, atomic images 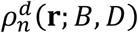 of a limited resolution *D* have a large peak in the atomic center (Fig. 1), now these peaks are surrounded by Fourier ripples. We show below that even when the amplitude of the ripples decreases, modeling this central peak only (*e.g*., Lunin & Urzhumtsev, 1984; Mooij *et al*., 2006) is insufficient to reproduce the maps accurately. As a consequence, considering this effect being similar to that from the atomic disorder (Jacobi *et al*., 2017) could be inappropriate and a number of Fourier ripples should be included into calculations.

**Fig. 1.**
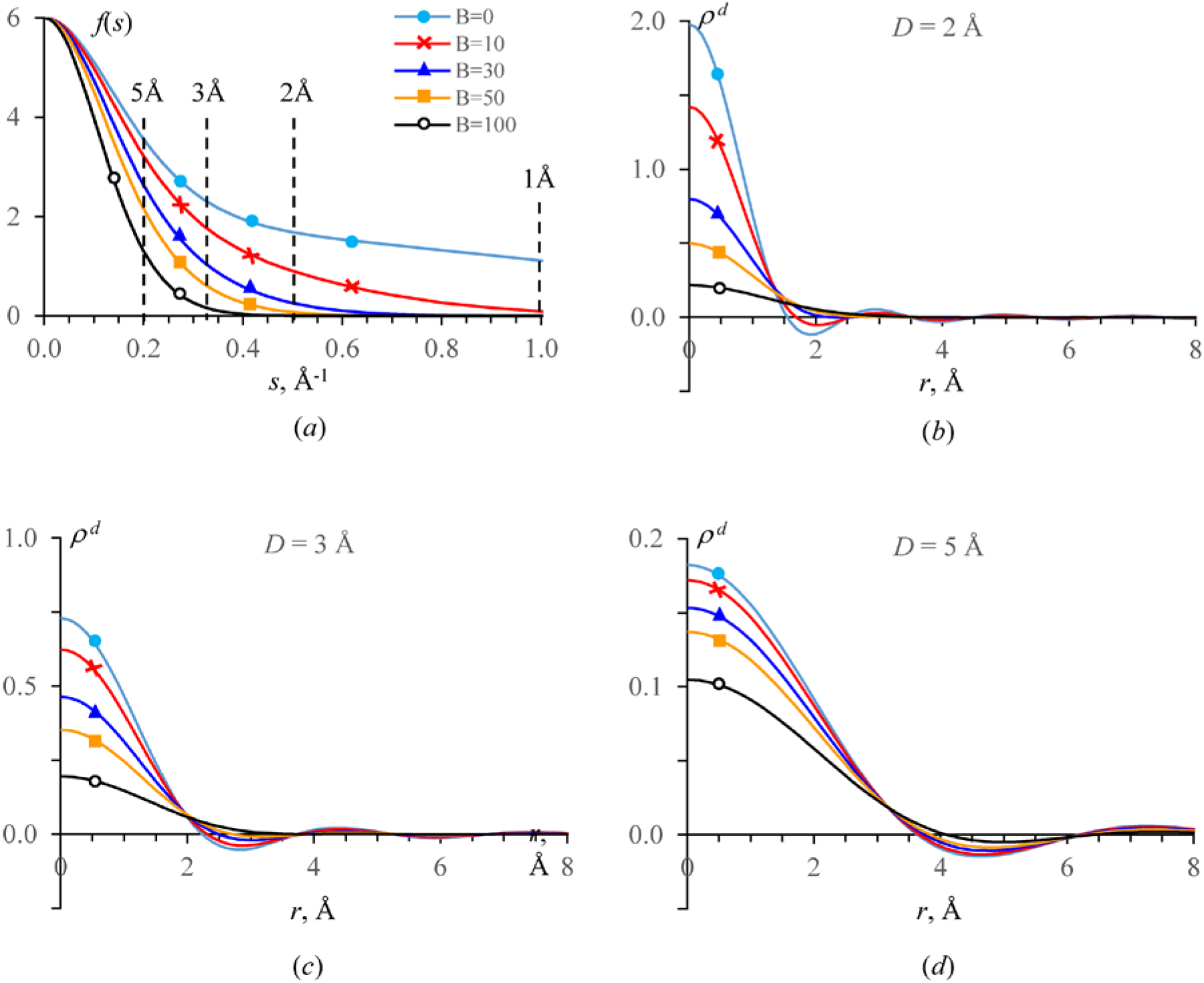
a) Scattering factor *f*(*s*) of carbon atoms with different *B* values. b, c, d) The radial component 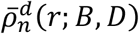 of the image of the respective carbon atoms calculated with a different resolution cut-off *D*. The scale for the image values is different in different figures. The figures use same color and marker code as defined in a).

Using atomic images with a truncation distance *r_max_* means that the direct map becomes a sum

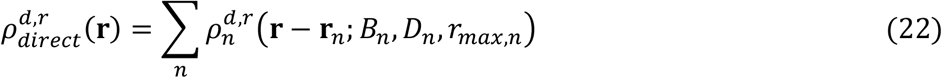

of truncated atomic images:

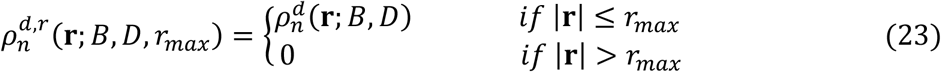

Unless the opposite is required explicitly, below we consider *r_max,n_* = *r_max_*, *D_n_* = *D*, the same values for all atoms. To simplify expressions, we omit *r_max_* when this does not lead to confusing.

### 2.2. Truncation distance and integral amount of the density

To choose the atomic truncation distance when calculating the density distribution, one may compare the amount of electron density *Z*(*r_max_*) inside the sphere of the respective radius *r_max_* with the full number of electrons *Z*(∞) of this atom (*e.g*., Agarwal, 1978). Divergence of these values could be an indicator of the map inaccuracy.

We start from a similar comparison for an image of an isolated atom. Since the atomic image at any given resolution is a spherically symmetric functions, we express the respective integrals via the radial component of the image, 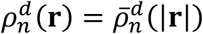:

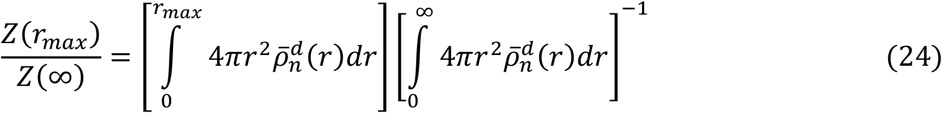

For an image of a point atom (14–15) this integral can be calculated in the closed form showing that the result does not rapidly converged, as it is often believed, but is an oscillating function (Appendix A)

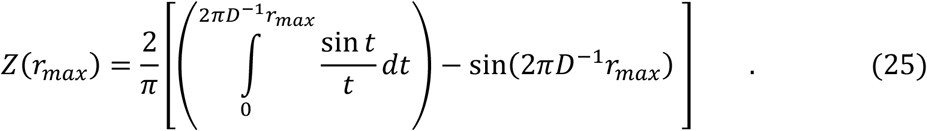

The first term in the brackets is the Sine integral function that tends to one for *r_max_* → ∞ (Fig. 2), while the second term is a sine function with the period *D*^−1^*r_max_*. The ripple amplitude decreases as approximately 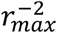 for the interference function (15), while the volume of such ripples in space is roughly proportional to 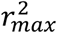 explaining why *Z*(*r_max_*) does not converge to *Z*(∞) and does not converge at all.

**Fig. 2.**
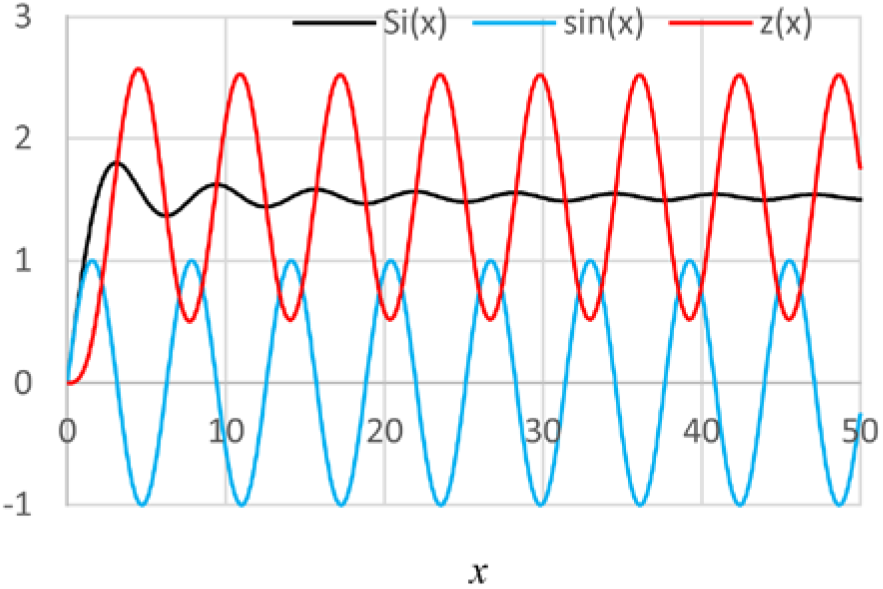
Functions describing the amount of density inside the sphere of the radius *x* for a point scatterer; equation (25). Sine integral Si(*x*), sin(*x*) and their unscaled difference z(*x*) = Si(*x*) – sin(*x*).

For a Gaussian atom 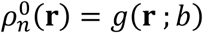 with *Z*(∞) = 1, the function *Z*(*r_max_*), calculated this time numerically, shows a similar oscillating behavior (Fig. 3) while the amplitude of oscillations of (24) decreases with increasing *b*. Previously, Tickle (2012) also mentioned a slow convergence of such kind of integrals. Like for the point atom, *Z*(*r_max_*) ≈ *Z*(∞) if *D*^−1^*r_max_* is integer or half-integer (Appendix A and Fig. 3). This condition is important for accurate map calculations, and the choice of the truncation distance as

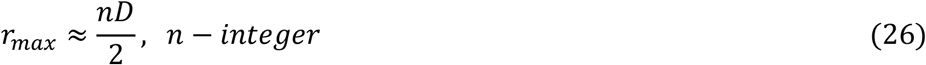

may improve map accuracy.

**Fig. 3.**
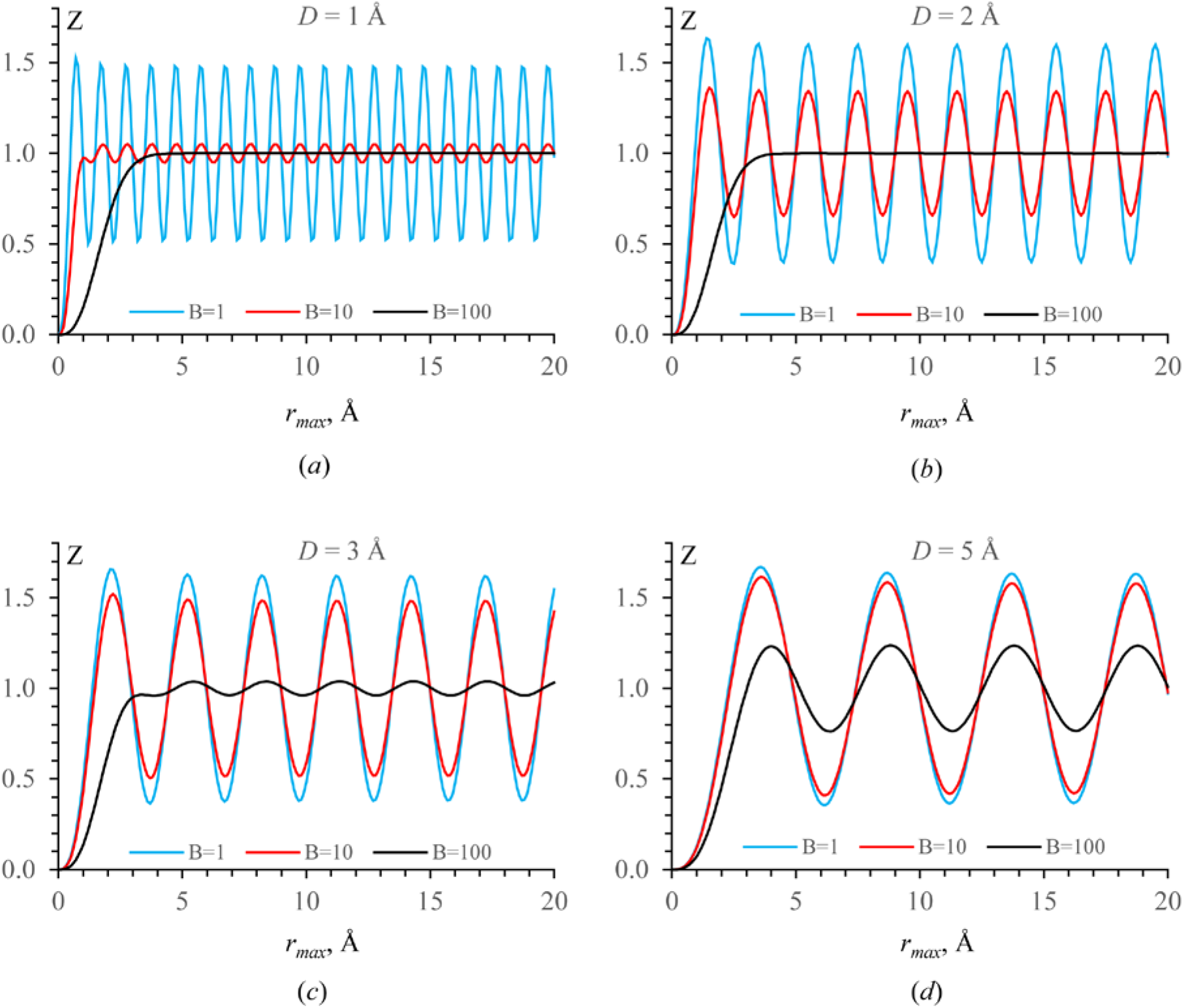
Integral amount of electron density *Z*(*r_max_*), as a function of the truncation distance *r_max_*, of an image of a Gaussian virtual atom; images are calculated with a different resolution cut-off *D* and for different *B* values.

### 2.3. Truncation distance and map values

Previous tests show that the truncation radius of the atomic image, *i.e*., the ‘tails’ of these images, significantly and non-monotonous influences the amount of the scattering density when calculating a model map. Next, we check how strong the value of density in a model map is affected by the ‘tails’ of images of neighboring atoms, depending on the number of these atoms and the distance to them, or, in other words, on the truncation radius of atomic images. We start from a simplified test when only the translationally equivalent atoms are considered as neighbors; we continue then with more realistic atomic models. Let us consider first a virtual cubic crystal with the cell parameters *a* = *b* = *c* = 10 Å in space group P1, with a single carbon atom with *B* = 10 Å^2^ in the origin. The Fourier map calculated as (11) at the resolution *D* = 3 Å had its value in the atomic center 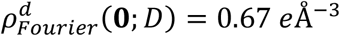. As discussed in Introduction, this value is the sum of contributions from an infinite number of carbon atoms, in origin of all unit cells of such infinite virtual crystal. To reproduce this density value in the directly calculated map 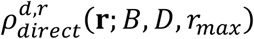, all atomic contributions should be included with a very large truncation distance, *r_max_* → ∞. Independently, we calculated the image of an isolated carbon atom using Fourier transform

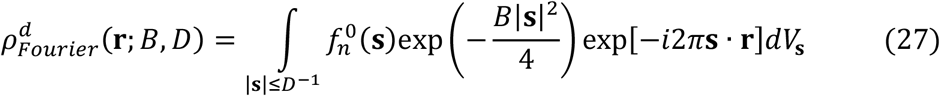

with *B* = 10 Å^2^ at the resolution *D* = 3 Å. Here 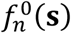 is the scattering function of the immobile carbon atom. In the center of the atom, 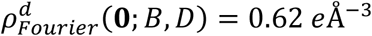. The difference with the Fourier map value, close to 8% and not negligible for refinement, is due to the contribution of ripples from neighboring carbons. In the second calculation, this contribution is missed, that is equivalent to calculating the direct model map 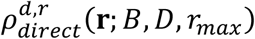 with the truncation distance *r_max_* <10 Å. Fig. 4 shows the value 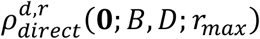 calculated for intermediate *r_max_* values. It approaches to 0.67 *e*Å^−3^ not monotonously with *r_max_* and much slower than one would expect. The reason for this is the same as above: the number on atoms at a distance *r* in roughly proportional to *r*^2^ while the ripples amplitude decreases as *r*^−2^.

**Fig. 4.**
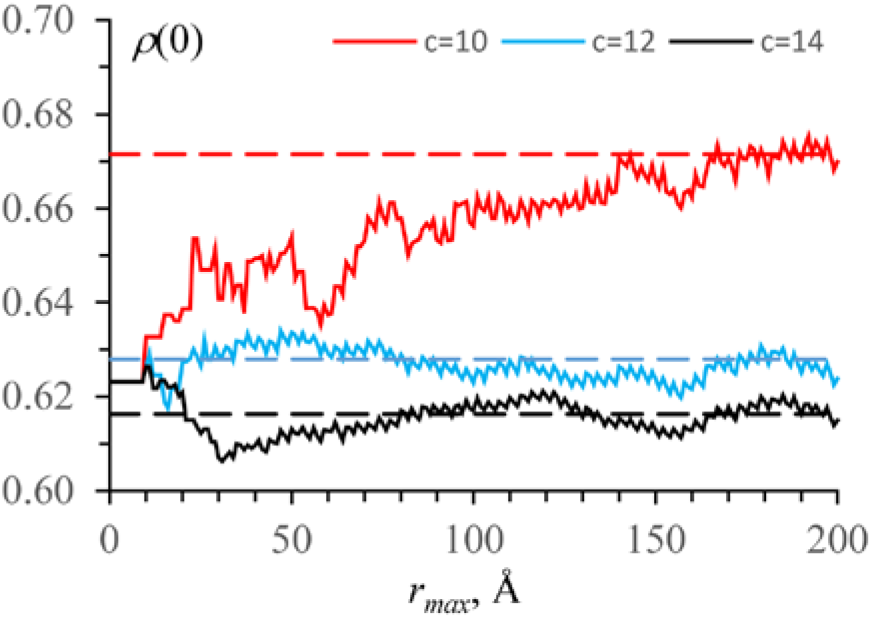
Map value 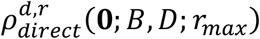 as a function of the truncation distance *r_max_*. The map of 3 Å resolution was calculated for a model composed of a single carbon atom in the origin with *B* = 10 Å^2^. Three virtual crystals were tested with unit cell parameters: *a* = 10 Å and with *b* = *c* = 10 Å, or *b* = 11 Å, *c* = 12 Å, or *b* = 12 Å, *c* = 14 Å. Broken lines show the respective 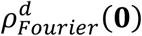 values.

This behavior indicates also one more feature of this test: in the cubic cell the contributions from neighboring atoms come ‘in phase’. In similar tests with orthogonal but non-cubic unit cells, *a* = 10 Å, *b* = 11 Å, *c* = 12 Å or *a* = 10 Å, *b* = 12 Å, *c* = 14 Å, the convergence was achieved at a much shorter distance, which stays significant. Note that the control value in the origin, 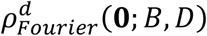, varies with the unit cell parameters.

### 2.4. Macromolecular maps and their comparison

Macromolecular crystals contain many atoms and have sufficiently large unit cells being quite rare cubic. The latter results let us believe that for such crystals a contribution of atoms from not neighboring unit cells is small and that different kinds of ‘special effects’ are hardly expected. On the other hand, they indicate that using only the central peak of atomic images may result in significant map errors.

To check these hypotheses for a protein model (Appendix B), we calculated various direct maps 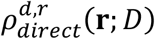, of a homogeneous resolution *D_n_* = *D* with the same truncation distance for all atoms *r_max,n_* = *r_max_*. We compare these maps with the Fourier map 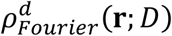 considered as the control one:

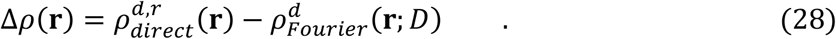

The considerations above show that a point-by-point map comparison

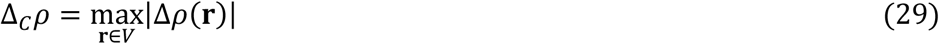

may be more informative than the traditional integrative metrics like the least-squares one

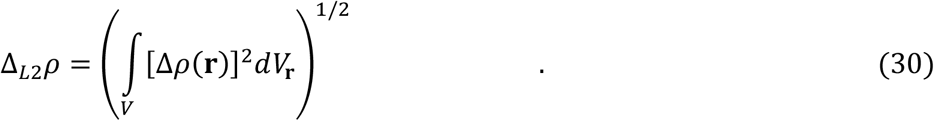

We calculate also the normalized difference

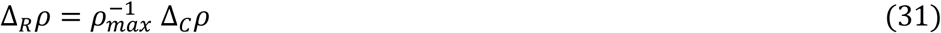

where

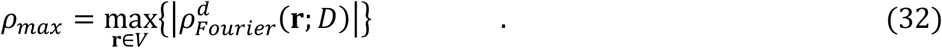

### 2.5. Protein model of identical atoms

In the first series of tests, all atoms of the model were assigned to be carbons with the same *B_n_* = *B* taken equal to 10, 30 or 100 Å^2^, changing from one test to another. The 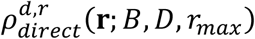 maps were calculated with the exact images of such atoms calculated as (27) at the resolution *D* equal to 2, 3 or 5 Å using X-ray scattering functions and cut then at some distance *r_max_* (23). These maps were compared with the respective Fourier maps 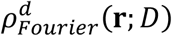 considered as the true answer. For these tests, the single source of the difference between the two maps was the truncation distance *r_max_* of the atomic contributions. Figs. 5a-c show Δ*_R_ρ* as a function of *r_max_* for different combinations of parameters *B* and *D*. As expected, this function is not monotonous. Agreeing with the previous considerations, smaller errors are observed for *r_max_* = *nD*/2 where *n* is an integer. The relative map discrepancy *Δ_R_ρ* grows with resolution value *D* at each *r_max_*. For *B* = 100 Å^2^, there is practically no difference between the curves at the resolutions *D* of 2 Å and 3 Å; this is expected since the difference between the respective scattering curves is small (Fig. 1a).

**Fig. 5.**
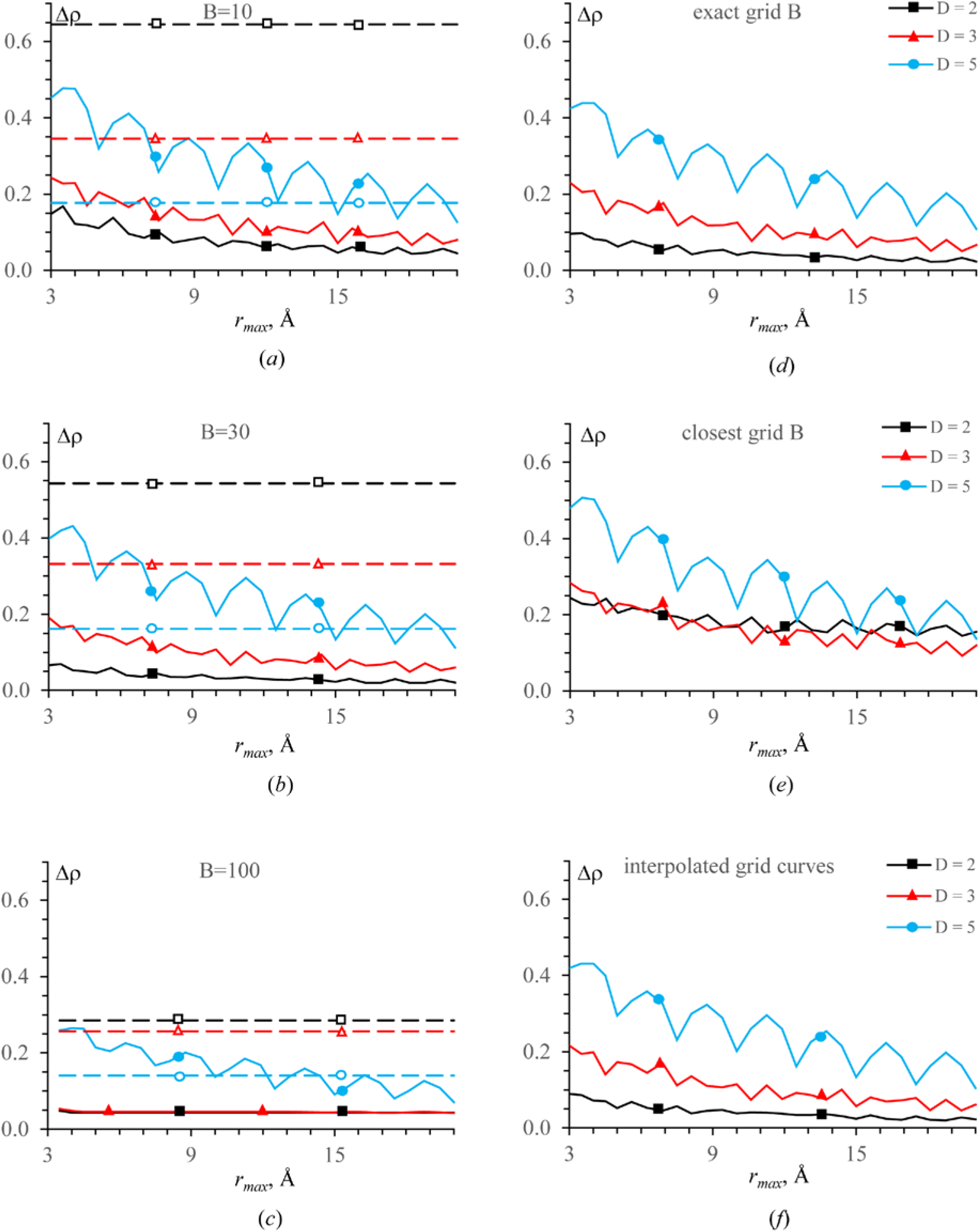
Maximal relative normalized difference Δ*_R_ρ*, in eÅ^−3^, as a function of the truncation distance *r_max_* for the maps calculated at different resolution values *D*. All atoms are considered as carbons. Control 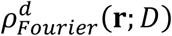 maps were calculated: a)-d) with the *B_n_* values corresponding to those in the respective model; e-f) 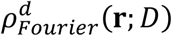 with the *B_n_* values taken from the PDB. Direct 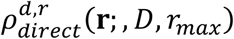 maps: a)-c) all atoms with the same *B_n_* value indicated in the plots; d-e) atomic *B_n_* replaced by the closest value from the *B*-grid; f) the curves for the closest *B*-grid values were interpolated. Broken lines show the value of the image of a carbon atom in its center also normalized by *ρ_max_*.

The average map difference normalized by *ρ_max_* (not shown) also oscillates with *r_max_*. However, the amplitude of oscillations is small, of order of 0.01 for the maps of 2 Å and 3 Å and of order of 0.02 - 0.04 for the maps at 5 Å. Thus, differently from *Δ_R_ρ*, this integral metric does not indicate existence of significant map discrepancies in particular points.

### 2.6. Protein models of carbons with different B

Identical *B_n_* values for all atoms may be seen as another source of confusion when calculating a model map. To evaluate this effect, in the second series of tests, we took the same macromolecular model with all atoms considered as carbon while varying *B_n_* values from one atom to another. Since it is computationally impractical to calculate an image for each atom with its own current *B_n_* using (27), we calculated a set of carbon images 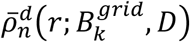 for twelve different 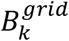 values, *k* = 1, 2, …, 12 (*B*-grid) in the interval from 5 to 100 Å^2^ (Appendix B).

First, we substituted each atomic *B_n_* by the closest value from the *B*-grid and used this model to calculated both 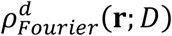 and 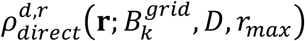 for the same three resolutions *D* as above. As in the previous test, the only source of errors was the truncation distance of the atomic contributions. The maximal deviation Δ*_C_ρ* did not change its shape (Fig. 5d). The respective curves at all three resolutions were between those for atoms with *B* = 10 Å^2^ and *B* = 30 Å^2^ corresponding to the minimal and mean *B* values in the model.

Moving toward a real situation, we kept the previous 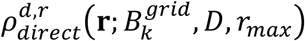 maps but calculate the control maps 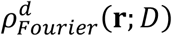 with *B_n_* values as they are in the PDB file; yet all atoms were considered as carbons. This added the second source of difference between the maps, namely inexact *B* values for the direct maps. Expectedly, the discrepancy Δ*_C_ρ* increased, especially at the highest resolution, *D* = 2 Å (Fig. 5e).

We repeated the last test getting an improved map 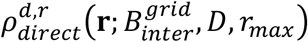. Now for each *B_n_* value we used the curve 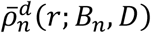 obtained by a linear interpolation of the curves 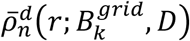 for the closest grid values, 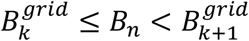, similar to DiMaio *et al*. (2015). This reduced the map error back to the values as they were in the initial test (Fig. 5f).

### 2.7. Models of mixed atomic types

The X-ray scattering functions of the principal macromolecular atoms (C, O, N, S) differ, at the first approximation, by a scale factor reflecting the number of electrons *Z_A_* in the atom of type *A* (Harker & Kasper, 1948). The direct map calculations can be simplified if the images of all atoms can be considered as, *e.g*., the carbon atom scaled by the number of electrons, *Z_A_*: *Z_c_* (Figs. 6a and 6b). Similarly, other types of atomic density, such as electron scattering potentials, can be rescaled to each other. The situation is different in cryoEM for neutral and charged atoms (Marques *et al*., 2019) for which this section is irrelevant.

**Fig. 6.**
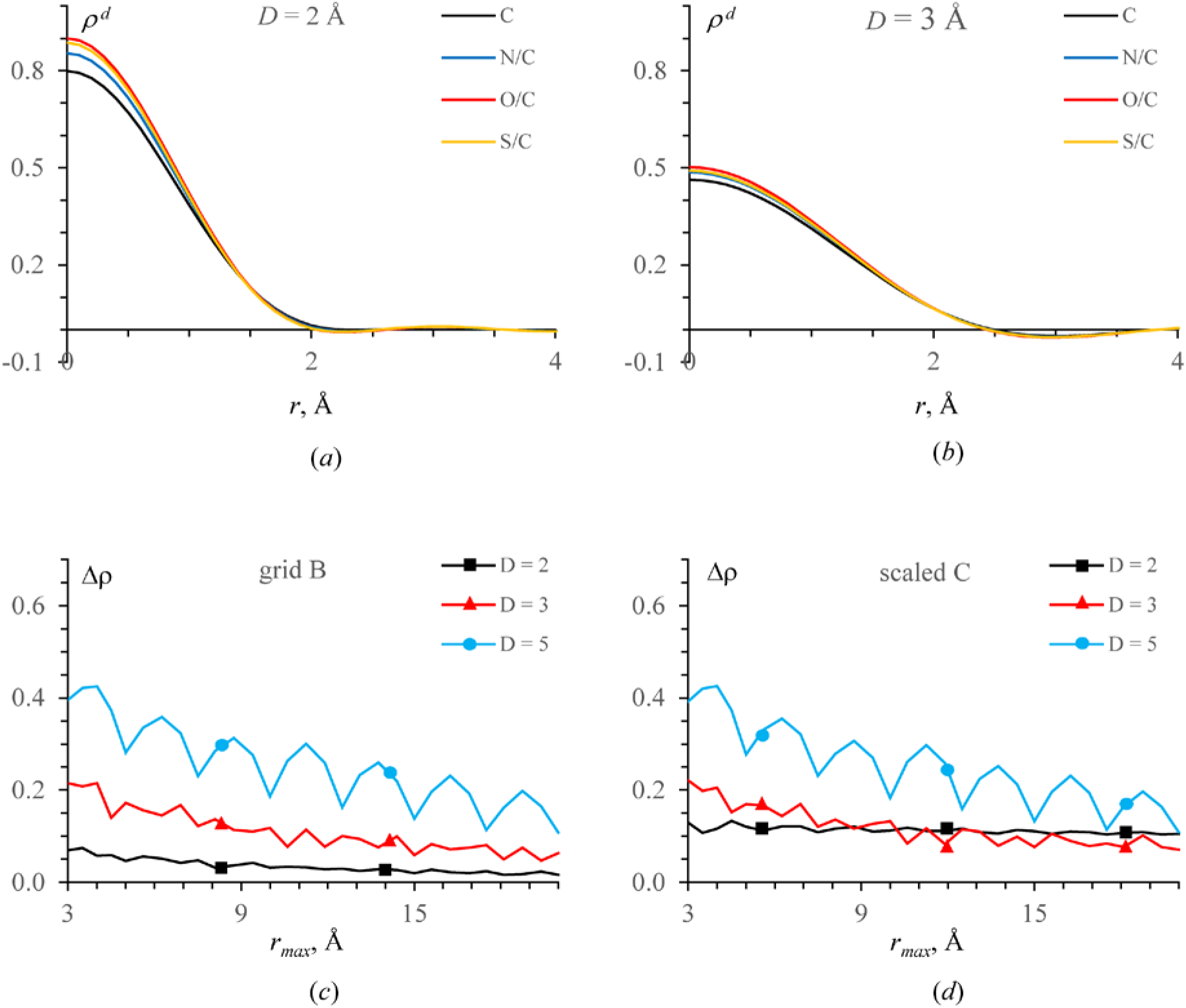
Maximal relative difference Δ*_R_ρ*, in eÅ^−3^, as a function of the truncation distance *r_max_* and calculated at the respective resolution *D*. a,b) Images of atoms with *B* = 30 Å^2^ at *D* = 2 Å and at *D* = 3 Å scaled by the number of electrons to the image of carbon. c) For both direct and Fourier maps, *B_n_* value of each atom was replaced by the closest value from the *B*–grid. All types of atoms were kept as they are in PDB. d) 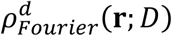 was conserved from the previous test; the carbon images rescaled by the number of electrons were taken to calculate 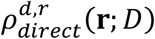.

First, we considered atomic types as they are in the model replacing each *B_n_* value by the closest one from the *B*-grid. We used this model to calculate both 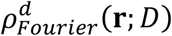 and 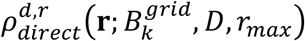 where the atomic images were prepared and used for each atomic type (C, N, O and S). As in the first tests, the single source of the difference between the maps was the truncation distance *r_max_*. Expectedly, introducing mixed atomic types practically did not change the previous results (Fig. 6c).

Then, keeping the *B* values from the previous test, the curves 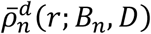 for all types of atoms were taken as those for carbon rescaled by the number of electrons. This practically did not affect the maximal deviation in the map of the resolution of 5 Å, slightly increased the error at 3 Å while increased it significantly at 2 Å.

### 2.8. PDB protein model

Finally, we simulated a more realistic situation. We calculated structure factors from the atomic model (Appendix B) with the parameters as they are in the PDB. With these structure factors, we calculated 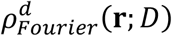 at the resolution of 2, 3 and 5 Å, as previously, considering them as the control maps. Independently, we calculated the images 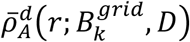 of atoms of the types *A* (C, N, O and S) at these three resolutions and for the *B* values from the same *B*-grid as above, 4 × 3 × 12 curves in total.

At each resolution *D* we calculated 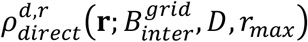 as a sum of atomic contributions of respecting atomic types. For each atom, its image was calculated by a linear interpolation of the ‘grid images’ for the grid values 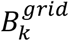 closest to the atomic *B_n_*. The maximal map discrepancy (Fig. 7a) is close to that from the last simplified test (Fig. 6c).

**Fig. 7.**
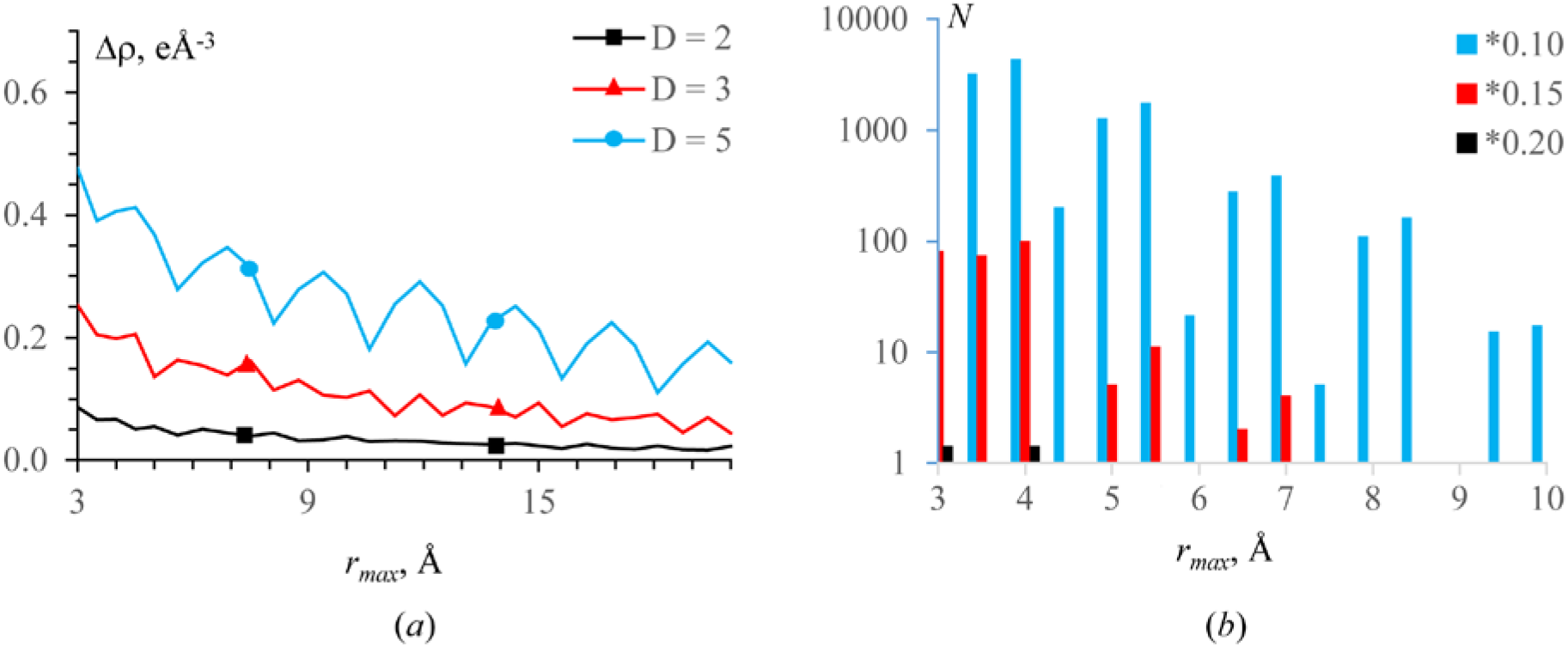
Map analysis for the test protein model. a) Maximal relative difference Δ*_R_ρ* as a function of the truncation distance *r_max_* and calculated at the respective resolution 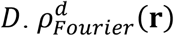 was calculated with the PDB values; the correct atomic types and the interpolated curves from the *B* –grid were used to calculate 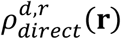. b) Number of grid nodes with the relative map difference Δ*_C_ρ* above a given part of the maximal 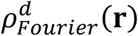 value (0.10, 0.15, 0.20, respectively); the numbers are given, as a function of the truncation distance *r_max_*, for the maps calculated at the resolution *D* = 3 Å. The histogram is given in a logarithmic scale.

As an integral measure, we calculated the number of points with large map discrepancy. This value decreases with *r_max_* also oscillating with a step equal approximately to *D*/2. For example, for the maps at *D* = 3 Å, it suggests *r_max_* = 6 Å as a possible choice, or *r_max_* = 7.5 Å if a higher accuracy is required (Fig. 7b). The peaks of the difference maps are distributed uniformly over the molecular region with no particular preferences.

Finally, we analyzed some maps visually. Fig. 8 gives an example of the maps calculated at the resolution of *D* = 3 Å. The 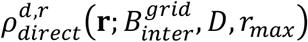 maps both with *r_max_* = 4 Å and with *r_max_* = 6 Å are in overall similar to 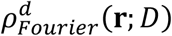. However, in a few regions, the map calculated with *r_max_* = 4 Å is poor (Fig. 8). The map 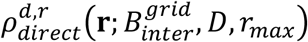 calculated with *r_max_* = 6 Å has no such imperfections. More tests are described in § 3.4.

**Fig. 8.**
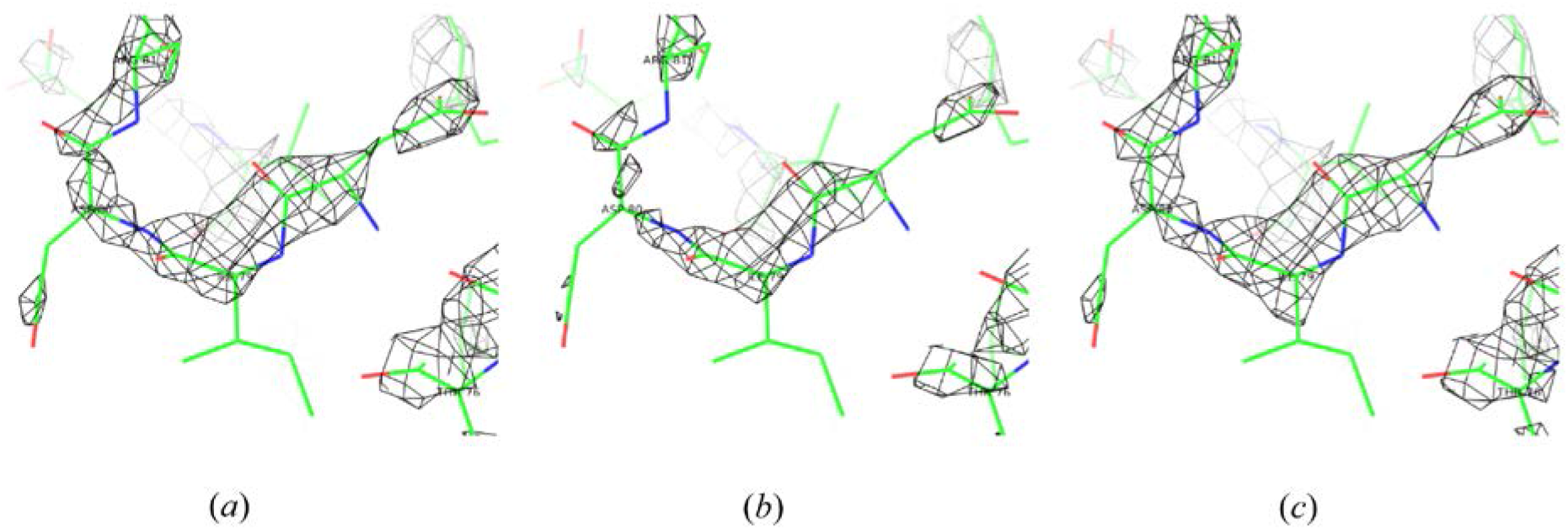
Maps for the test protein model. Map are calculated at the resolution *D* = 3 Å with the correct atomic types and *B* values as they are in PDB. Interpolation curves are used for the direct map calculation. a) Fourier map 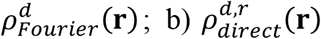 with *r_max_* = 4 Å; c) 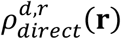 with *r_max_* = 6 Å. All maps are contoured at 1σ level.

### 2.9. Practical questions of model map calculations

After the first macromolecular refinement program (Diamond, 1971), real-space refinement in macromolecular crystallography has been substituted by reciprocal-space refinement starting end of 70ths. Being limited for a while by its use in computer graphics programs, *e.g., coot* (Emsley & Cowtan, 2004), real-space refinement got a new interest with the recent progress of cryo EM (Chapman *et al*., 2013; Murshudov, 2016; Afonine, Poon *et al*., 2018). M

A model map 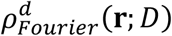 for comparison with the experimental one can be calculated by the inverse Fourier transform with the model structure factors. However, the fastest way to get these factors is another Fourier transform, that of a model density calculated with atomic radii in a fine enough grid. A larger grid step can used by introducing an artificial extra atomic displacement parameter which in turn needs larger atomic radii (Ten Eyck, 1977; Navaza, 2002).

The tests above show that a direct map composed of the individual atomic images can be calculated instead avoiding Fourier transforms at all. However, the truncation distance of atomic images should be larger than the traditional values used for atomic densities (2.5-3 Å, as a rule). Increasing this distance improves 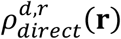 but increases the CPU time.

At the same time, the direct map calculation solves another important problem, not seen previously in crystallographic studies. Due to “resolution revolution”, real-space refinement of atomic models *versus* cryo EM maps, resolution of which varies from one molecular region to another, becomes more and more critical. Such inhomogeneous-resolution model maps can hardly be calculated as the Fourier ones.

Below we propose a method to calculate atomic images of a given resolution as analytic functions of all atomic parameters up to any truncation distance. Such analytic expression allows accurate calculation of model maps and, therefore, an efficient refinement of atomic parameters. Using the suggested approach, the local resolution becomes a parameter of these analytic functions. With this development, one does not need any more to prepare atomic images at a particular resolution, but complete the list of parameters, such as atomic coordinates and displacement factor, by the local resolution and refine it together with these parameters.

## 3. Analytic approximation to an atomic image

### 3.1. Ω-function and its features

Atomic distributions, or rather respective scattering functions of immobile atoms, are traditionally modeled by a linear combination of a few three-dimensional origin-centered Gaussian functions:

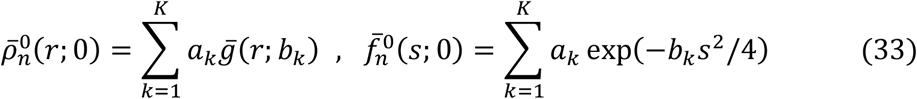

(*e.g*., Grosse-Kunstleve *et al*., 2004). A Gaussian function has an important feature that its convolution with another Gaussian still be a Gaussian but with the modified value of its parameter:

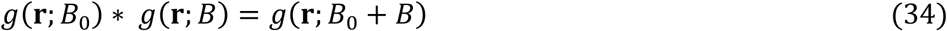

In other words, knowing the atomic density for an immobile atom, with *B* = 0, this gives such density for an atom with a non-zero *B* value:

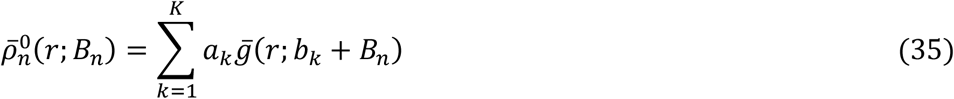

As previously, an overline indicates the radial component of a function spherically symmetric in space.

We would like to have a similar approach to atomic images at a limited resolution. When ignoring the Fourier ripples, one can model the peak in the origin also by a Gaussian (*e.g*., Lunin & Urzhumtsev, 1984; Sorzano *et al*., 2015, and references therein) or by a polynomial (Mooij *et al*., 2015). However, the tests above show that modeling the central peak only is insufficient to obtain accurate 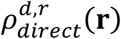 maps by a sum of such contributions. Chapman (1995, 2013) suggested to express atomic images using a step-function approximation to the scattering factors up to the chosen resolution limit. Alternatively, for an image of an immobile atom, *B_n_* = 0, of each required type at a given resolution, Urzhumtsev & Lunin (2022) suggested to decompose it into a sum of terms

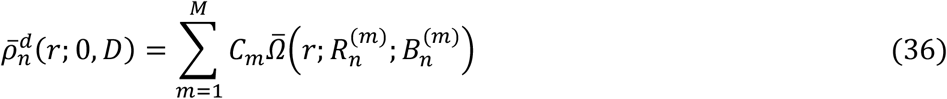

expressed through the specially designed function

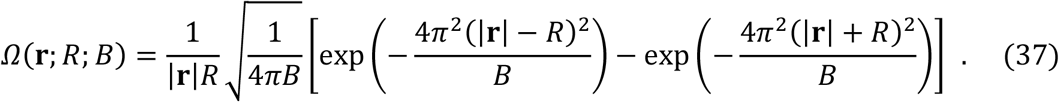

This function describes an image of a unit charge uniformly distributed over a spherical surface of the radius *R* with the isotropic disorder described by the Gaussian distribution with the parameter *B*. It represents a kind of a solitary spherical wave (Fig. 9) when *R*^2^ > 3*B*/(8*π*^2^) and a peak in the origin otherwise. It approaches the Gaussian function (18) when *R* → 0. Another important feature of *Ω*(**r**; *R, B*), which directly follows from its definition, is that its convolution with a Gaussian does not change it form but only the value of its parameter *B*_0_:

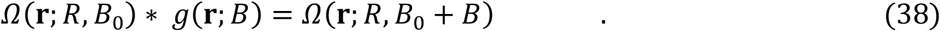

**Fig. 9.**
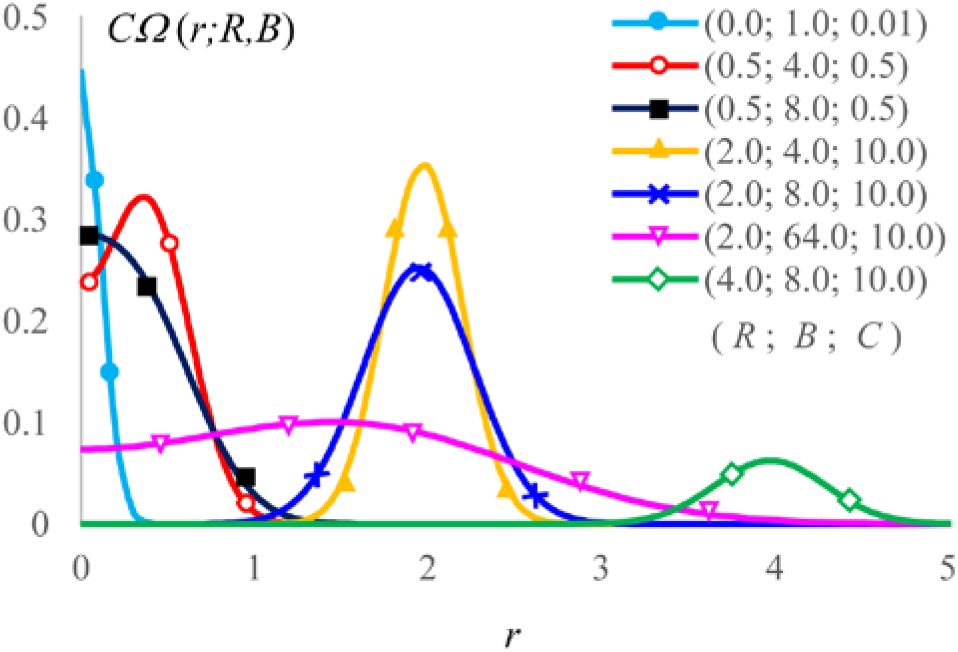
**Function** *CΩ*(**r**; *R; B*). The radial component of the function is shown for several values of its parameters (*R; B; C*).

### 3.2. Shell decomposition of atomic images

Decomposition (36) of 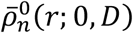 images of immobile atoms at a given resolution into a sum of *Ω*-functions is computationally similar to a decomposition of the atomic densities into a sum of Gaussian functions. In both cases, for a given and once calculated set of the decomposition coefficients, it gives automatically the respective curve for an atom with any *B:*

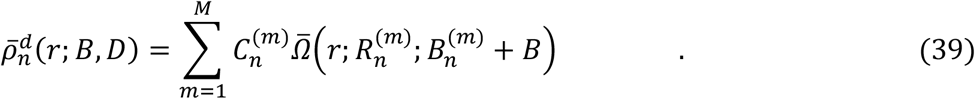

We call it the shell decomposition of the atomic image 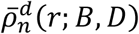. The term with 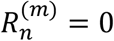 is a Gaussian describing the peak in the origin; eventually (39) may contain a few such terms. Other terms describe the distribution in spherical shells. The number *M* of terms in (39) is defined by the required truncation distance *r_max_* and the accuracy. Interference function (15) and atomic images have one positive and one negative ripple approximately per distance *D*. Numeric tests show that modeling each ripple by a single term, *i.e*., with *M* ≈ 2*r_max_/D* terms for the chosen *r_max_*, gives typically the maximal error of order of 2·10^−4^ - 3·10^−4^ with respect to the value of the central peak. Fig. 10 illustrates that including a few terms with 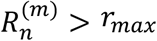 may improve the result because of a non-zero ‘width’ of the 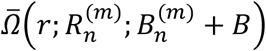 peak. An extrapolation (39) for terms with typical *B* values reduce this discrepancy by an order of magnitude further. If a higher accuracy required, extra terms can be added to model minor residual ripples.

**Fig. 10.**
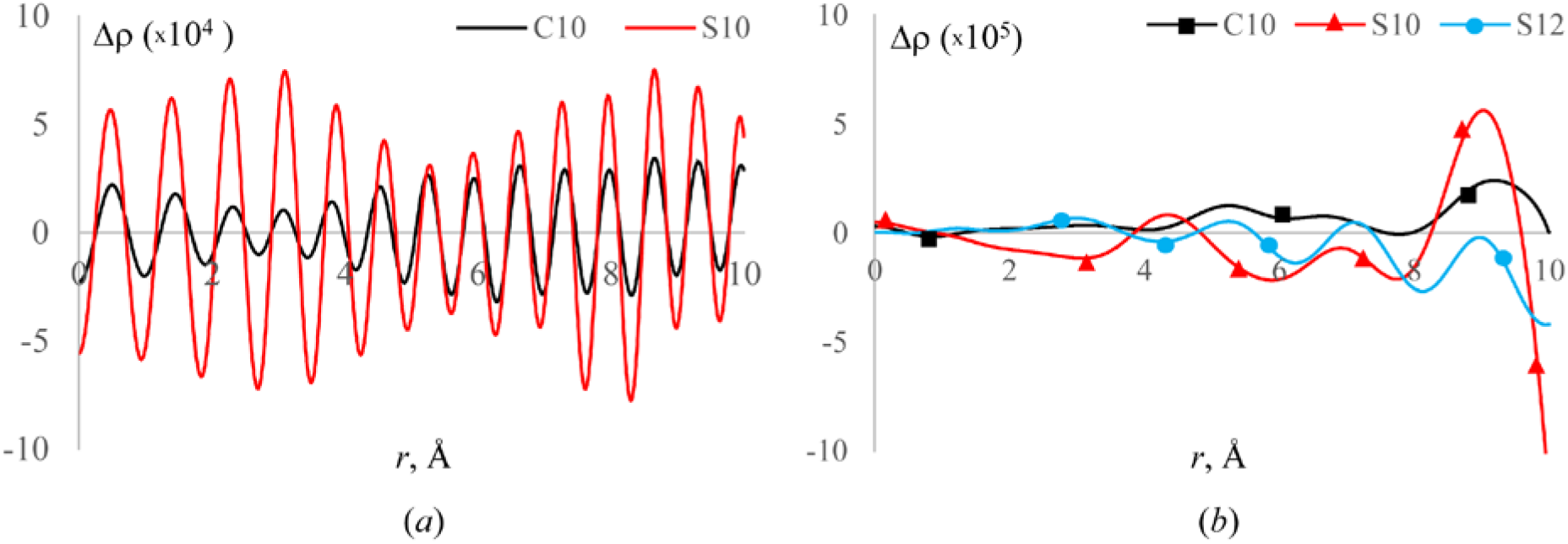
Discrepancy, in eÅ^−3^, between the images of C and S atoms at 2 Å resolution and their shell decomposition. a) Difference between the exact images of immobile atoms, *B* = 0 Å^2^, and their decomposition with *M* = 10 terms at the interval *r* ≤ 10 Å. b) Difference between the exact images of the atoms, *B* = 30 Å^2^, and the decomposition for immobile atoms with all *B*^(*m*)^ increased by *B* = 30 Å^2^ (39) and with *M* = 10 terms (curves C10 and S10). Addition of two terms for S, with *R*^(11)^ ≈ 11 Å and *R*^(12)^ ≈ 12 Å, improves the decomposition at the right bound of the interval (curve S12).

Parameters of the shell decomposition of an atomic image, or of any other oscillating function in space, are estimated from the shape and position of the Fourier ripples and refined then. If a higher accuracy is required, the decomposition can be completed by a new iteration applied to the residual curve. Details of the procedure are discussed in (Urzhumtseva *et al*., 2022).

### 3.3. Calculations of maps of inhomogeneous resolution

Crystallographic experimental maps are usually calculated as the discrete Fourier transform, in all grid points at a time, and supposed to have the same local resolution in all their regions. The situation is different for the maps of electrostatic scattering potential in cryo EM (Cardone *et al*., 2013). The variation of the local resolution is traditionally represented by a color map and is hardly available from structural data bases such as EMDB (Lawson *et al*., 2016) or EMPIAR (Iudin *et al*., 2016).

To calculate the respective model inhomogeneous-resolution maps, the resolution of each atomic image should fit the local resolution and is eventually could be refined in the same way as the displacement parameter. Presenting an atomic image as a double convolution (20) with *δ^d^*(**r**; *D*) defined by (14) suggests that the shell decomposition of

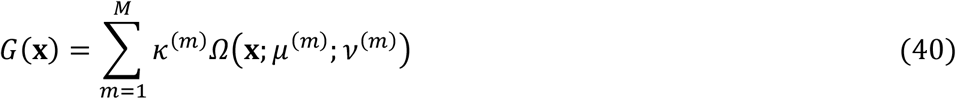

automatically gives an expression for an image of a Gaussian atom 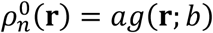 possessing of the displacement parameter *B* as an analytic function not only of the displacement parameter *B* but also of the resolution *D:*

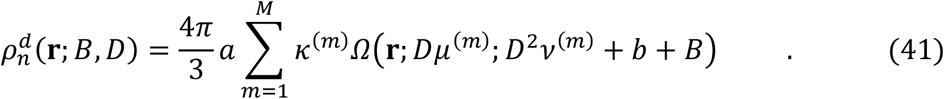

Here we use the rescaling property of the function *Ω*(**x**; *μ*, *v*) which follows immediately from its definition (37):

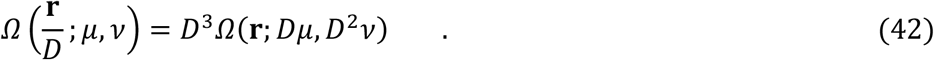

Finally, since an atomic density of an immobile atom can be represented by a sum of a few Gaussians, *K* ~ 2 – 5, the image of any atom at any resolution *D_n_* with any displacement factor value *B_n_* can be expresses as a sum (41) over these Gaussians with their own (*a; b*):

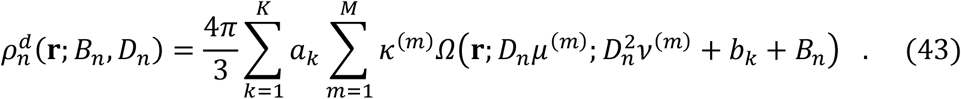

In this sum, all coefficients except *B_n_* and *D_n_* are tabulated once forever, *e.g*., Grosse-Kunstleve *et al*. (2004) for (*a_k_*, *b_k_*) and Lunin & Urzhumtsev (2022) for (*κ*^(*m*)^, *μ*^(*m*)^, *v*^(*m*)^). In other words, now the local resolution can be directly associated with the atom; we note this now by *D_n_* instead of *D*, common for all atoms, which is no longer correct. It characterizes the map against which the atomic position has been determined. This means automatically that this value can be refined together with other atomic parameters. This also means that the model map value in each point can be calculated considering the local map resolution, and this can be done in a single run, with no Fourier transforms applied.

Each of decompositions (36) and (43) has its own place and role in refinement of atomic models. The former has less terms and is appropriate at intermediate refinement stages when the resolution can be considered as a constant in the whole cell or region by region. Also, one can use it when a Gaussian approximation to scattering functions is not valid or when they are defined numerically (*e.g*., Fox *et al*., 1989; Brown *et al*., 2006; Sorzano *et al*., 2015; Murshudov, 2016). The latter decomposition allows refinement of a local resolution individual for each atom and is appropriate for final stages of the refinement, especially in cryo EM.

### 3.4. Map calculations using shell decomposition

#### 3.4.1. Data preparation

To validate the observations and conclusions obtained above with the test model, we made complementary tests with a model of IF2 solved previously at the resolution of 2 Å (Simonetti *et al*., 2013). The model composed of approximately 2900 atoms (C, N, O, S) was placed in the center of a virtual orthogonal unit cell in space group P1 with the size 80×120×100 Å. Model structure factors **F**_*model*_ (**s**) have been calculated at the resolution of 2 Å and the control Fourier maps *ρ_Fourier_*(**r**) were calculated as (6) at the resolution of 2, 3, 4 and 5 Å. To simplify the analysis, all maps were calculated in the same grid 120×180×150. We calculated the images of immobile atoms of each type (see Fig. 1) and calculated their decomposition coefficients (36) up to the distance 10 Å, with one term per ripple, at the resolution of 2 Å and 3 Å; *M* = 11 and *M* = 8, respectively (Table 1). The maximal difference between the image and the decomposition was equal to 3·10^−4^ and 4·10^−4^, respectively (the peak values are 1.98 and 0.73 eÅ^−3^)

**Table 1.**
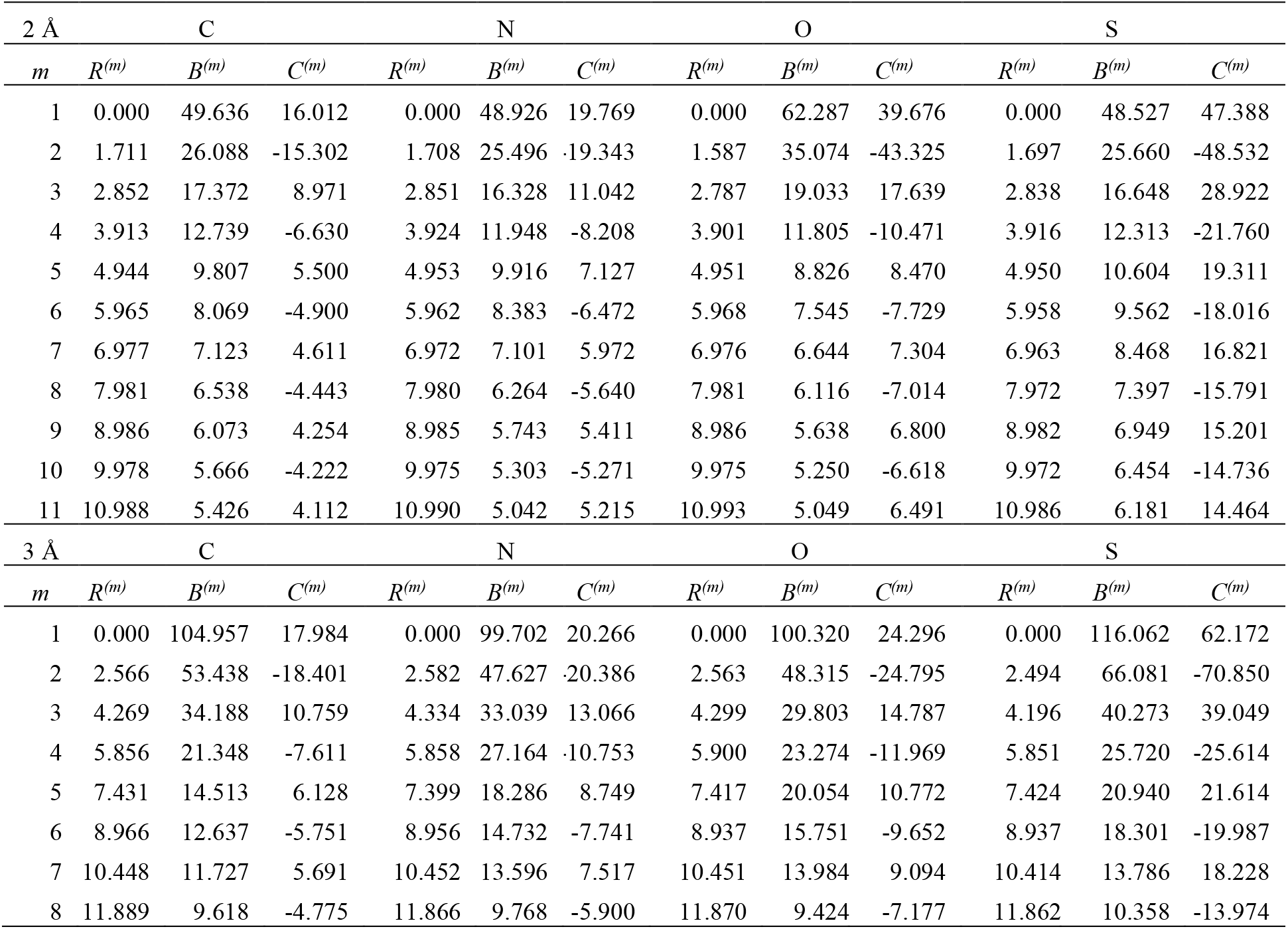
Coefficients of the shell decomposition of several atomic images. The decomposition was done up to the truncation distance *r_max_* = 10 Å, at the resolution of 2 Å and 3 Å. Note the presence of several terms with *R*^(*m*)^ > *r_max_*.

#### 3.4.2. Map of a homogeneous resolution

First, we calculated the map (39) at the resolution of *D* = 3 Å with the truncation distance *r_max_* = *nD* equal to 9 Å, with *n* = 3. The total number of the grid nodes in the unit cell is 3,240,000 and about 4% of them have the value above 0.1 eÅ^−3^ which corresponds roughly to 1*σ* cut-off; this agrees with previous studies (Urzhumtsev *et al*., 2014).

The difference |Δ*ρ*(**r**)| between the direct and exact maps of the same resolution (28) was above 0.1 eÅ^−3^ in only 22 grid nodes with the maximal deviation 0.12 (at this resolution, the images of the carbon atom in its center have the respective value 0.73, 0.46 and 0.20 eÅ^−3^ for *B* = 0, 30,100 Å^2^).

With reducing *r_max_*, the number of outliers above 0.1 eÅ^−3^ increases near exponentially, and the maximal deviation near linearly. The map quality is yet sufficient with *r_max_* = 6.0 Å, *n* = 2, and approaches to critical with *r_max_* = 4.5 Å, with *n* = 1.5 (Fig. 11). The points with deviations above 0.1 are inside the core of the molecule while deviations in more disordered molecular regions are slightly below 0.1 eÅ^−3^ (Fig. 12). Taking next recommended radius, *r_max_* = 3.0 Å with *n* = 1, results in very large errors; for example, the number of grid nodes with the deviation above 0.1 eÅ^−3^ approaches to 200,000.

**Fig. 11.**
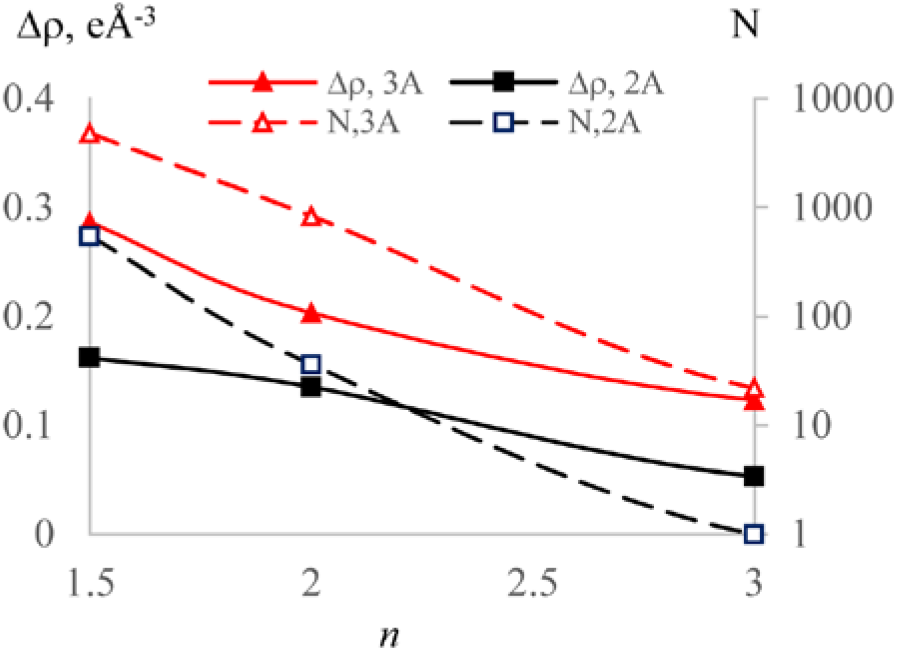
Comparison of the direct and Fourier maps for IF2. Solid lines show, as a function of *n* defining the truncation distance *r_max_* = *nD*, the maximal map deviation, Δ*ρ*, in eÅ^−3^; dotted lines show, in logarithmic scale, the number *N* of grid nodes with the discrepancy above 0.1 eÅ^−3^. Results shown are for the resolution *D* equal to 3 Å and to 2 Å.

**Fig. 12.**
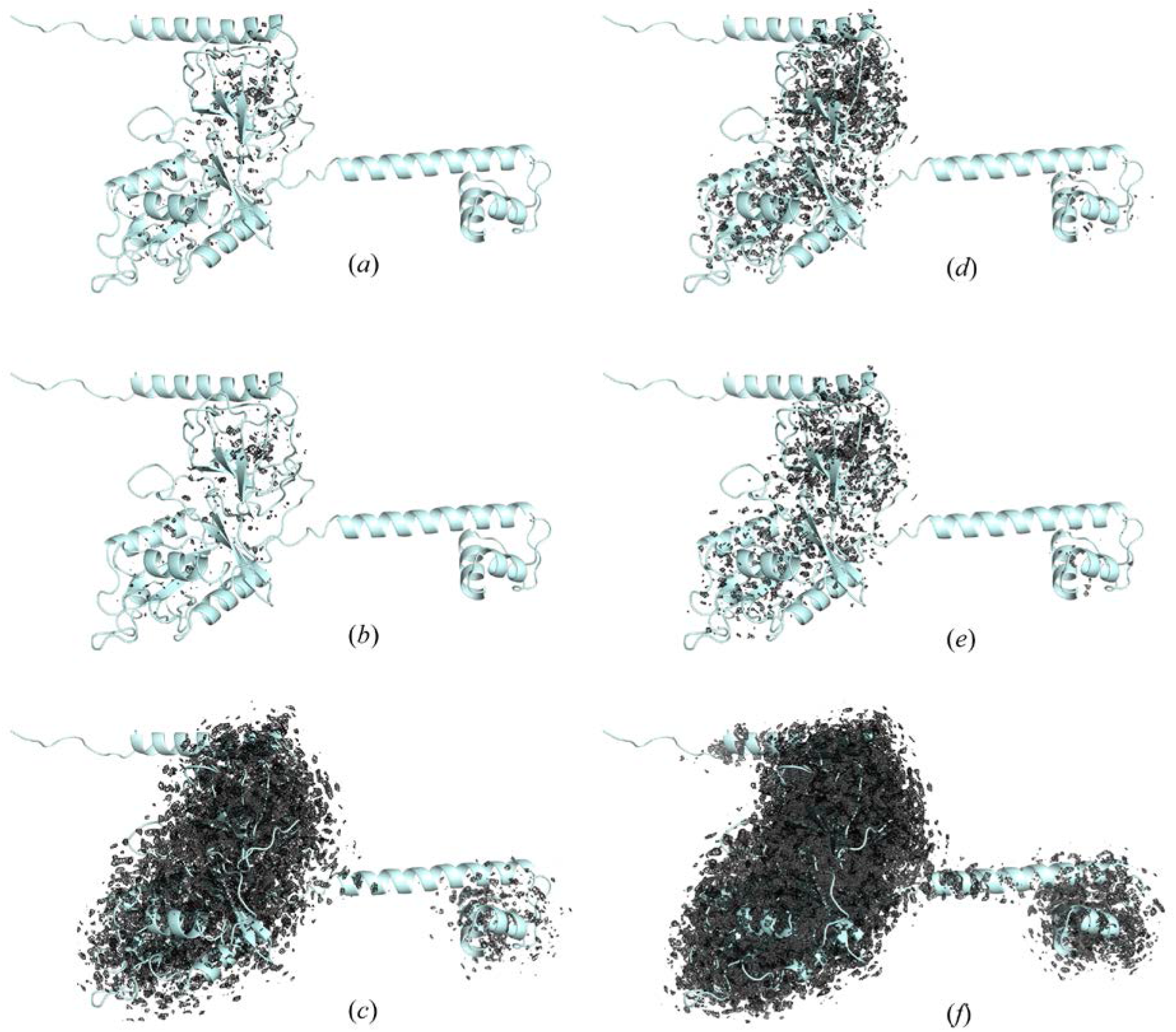
Absolute value of the difference between the direct and Fourier maps for IF2 at the resolution of 3 Å. The maps are shown for different combinations of the truncation distance *r_max_*, in Å, the number *M* of the decomposition terms, and the map cut-off *ρ_cut_*, in eÅ^−3^. Values of (*r_max_, M, ρ_cut_*) are: a) (6.0, 5, 0.10); b) (6.0,4,0.10); c) (6.0,4,0.05); d) (4.5,4,0.10); e) (4.5,3,0.10); f) (4.5,3,0.05).

One may expect that the terms in (39) with *R_m_* much larger than *r_max_* are insignificant. Indeed, when we removed all terms *m* >5 (corresponding to *R_m_* > 8.9 Å) for *r_max_* = 6.0 Å, or all terms *m* >4 (corresponding to *R_m_* > 7.3 Å) for *r_max_* = 4.5 Å, the maps changed only marginally. Removing next term (*R_m_* > 7.3 Å for *r_max_* = 6.0 Å and *R_m_* > 5.8 Å for *r_max_* = 4.5 Å) introduces some modifications still insignificant (Fig. 12) while removing one more term, with *R_m_* ≈ *r_max_*, completely breaks the maps (not shown).

We observed the same behavior for the calculations at the resolution of *D* = 2 Å with the truncation distance *r_max_* = *nD* equal to 6, 4, 3 and 2 Å. However, the maximal difference between the direct and exact maps and the number of outliers above 0.1 eÅ^−3^ decreased in comparison with those for *D* = 3 Å (Fig. 11). Moreover, at this resolution, the images of the carbon atom in its center have larger values, those of 1.98, 0.80 and 0.22 eÅ^−3^ for *B* = 0, 30,100 Å^2^ showing that the relative accuracy improves.

As a result of these tests, we conclude that for real-space refinement at the resolution *D*, the truncation distance *r_max_* recommended may be equal to 1.5*D* at earlier refinement stages and 2*D* at the end of refinement. Obviously, larger values such as *r_max_* = 2.5*D* as may be tried while they increase the computation time. This may be more important at lower resolution however this makes *r_max_* too large. When considering the decomposition with one term per ripple, the series (39) should include all terms up to *R_m_* ≈ *r_max_* plus the next one. Again, for final accurate calculations, it may be completed by one more following term.

#### 3.4.3. Map of an inhomogeneous resolution

To illustrate the calculation of inhomogeneous-resolution maps, the option crucial for cryo EM studies, for each atom of the IF2 model we assigned artificially a ‘local resolution’ *D_n_* increasing linearly from 2 Å in the central region of the model (sphere of the radius of 10 Å) to 5 Å at the periphery, beyond 40 Å from the model center. The truncation radius *r_max,n_* was taken as 2*D_n_*. Figs. 13a,b illustrate both sources of blurring the map; one can observe the peaks for less ordered atoms but at higher resolution (closer to the center) or, inversely, for a distant but more ordered atoms of the N-domain. Fig. 13c shows a continuous loss of resolution along the helix between two IF2 domains.

**Fig. 13.**
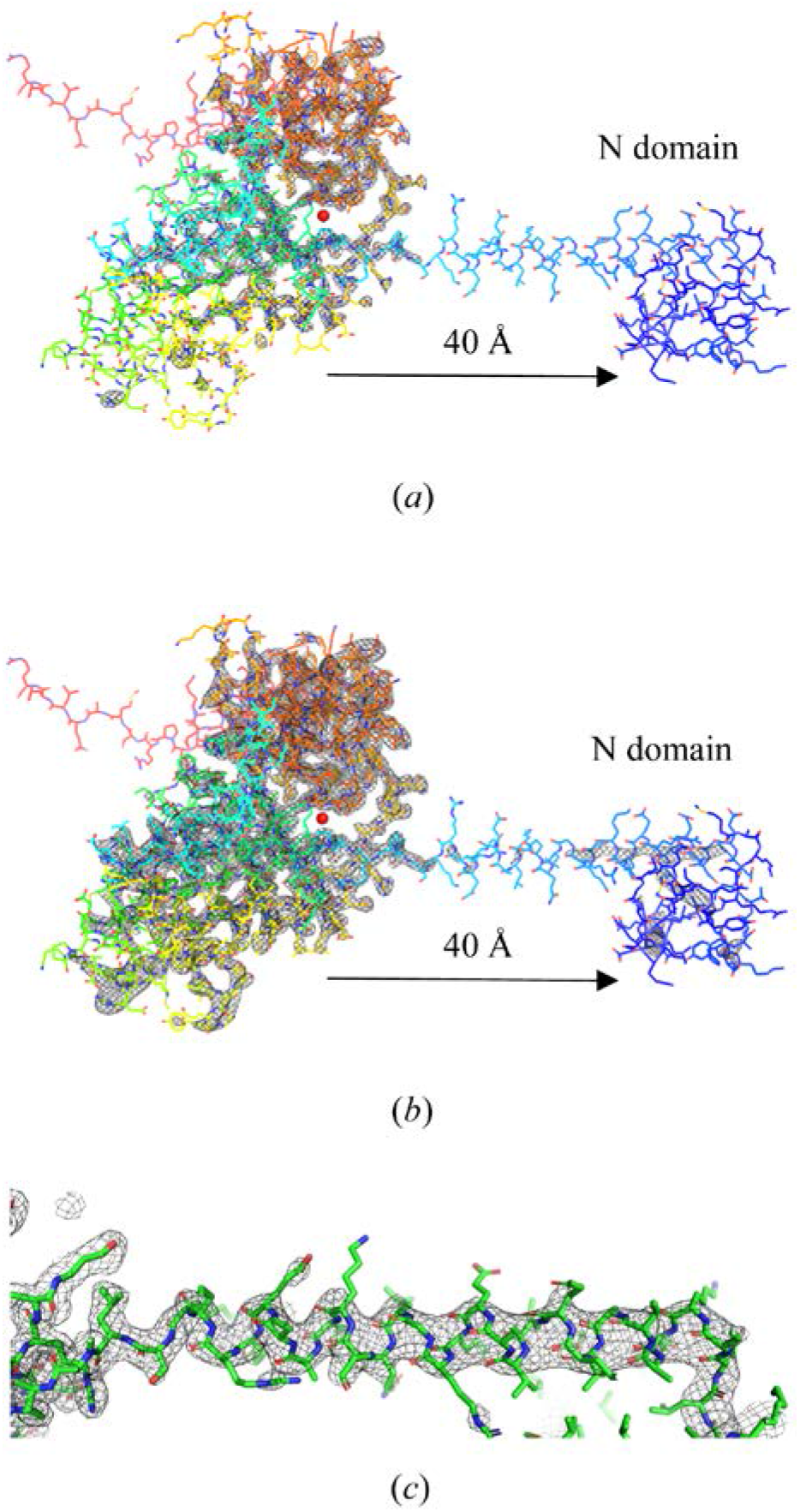
IF2 map of an inhomogeneous resolution calculated with the shell decomposition. a,b) Overall view; resolution varied from 2 Å in the model center to 5 Å at the model periphery. Map cut-off 1.0 eÅ^−3^ (a) and 0.75 eÅ^−3^ (b). Model is colored according to the *B* value. Model center is shown by a red dot. c) Loss of resolution with distance to the model center following the helix linking two domains (rotated by 90° with respect to the top images). Map cut-off 0.5 eÅ^−3^.

## 4. Discussion

Real-space refinement of atomic models improves these models by their fit to the experimentally obtained maps of an electron or nuclear density in crystallography, or of the scattering electrostatic potential in cryo EM. This procedure has several advantages with respect to reciprocal-space refinement, is complementary to it in crystallographic studies and is the principal procedure in cryo EM. A simplified procedure of a ‘fast-and-dirty’ refinement of atomic coordinates with point atoms (Afonine, Poon *et al*., 2018) is efficient at the first stages but leaves the model with some residual errors and is incapable to estimate atomic displacement parameters. An accurate real-space refinement of atomic models requires the model maps to mimic imperfections of the experimental ones. Such model maps can be calculated as a sum of atomic contributions (atomic images at given conditions) represented by oscillating functions in space. An analytic decomposition of these images using a specially designed function *Ω*(**r**; *R; B*), or *Ω*(**x**; *μ, v*) when expressed in dimensionless parameters, solves all principal problems. Below, we summarize some of them.

An image of an atom of any chemical type at any resolution and with any displacement factor can be represented analytically as a weighted sum of *Ω*(**x**; *μ, v*) terms. This decomposition contains both constant and variable parameters, to be refined. The first ones are calculated once forever for the image of the immobile atom of a given type and chosen accuracy. The second group includes the atomic displacement parameter and the resolution of the atomic image, as it is seen in the given map. In each space point, a model density map value becomes an analytic function of atomic parameters, *i.e*., their coordinates, displacement parameters and the local resolution values assigned now to every atom.

As a consequence, for a score function describing the model-to-experimental maps fit, its partial derivatives with respect to all atomic parameters become also analytic functions. All these parameters, including the local resolution, can be ‘real-space’ refined using gradient methods, with no need in structure factors and Fourier transform.

A model density map, even when its resolution varies from one its region to another, can be calculated in a single run, with no Fourier transform needed. The truncation distance and the number of terms can increase between initial refinement runs and the final ones to handle between coarser and faster calculations and those more accurate but taking more CPU time. Eventually, highly automated refinement programs like *Phenix* (Liebschner *et al*., 2019) may suggest automatic refinement protocols varying, as a function of data, the sets of refined parameters, atomic truncation distances and other parameters.

Being associated with atoms, the local resolution can be reported in the PDB files together with the coordinates and displacement factors; this value is a measure of confidence of the atomic parameters charactering the map region in which the atom has been identified.

For practical calculations it is important that, when the variation of the local resolution can be neglected, one can use a simplified form of a decomposition of images at a given resolution into a sum of *Ω*(**r**; *R, B);* this reduces the number of terms, in turn reducing CPU time and improving convergence; this may be the principal option at earlier and intermediate stages of refinement.

## Acknowledgement

AU acknowledges Instruct-ERIC and the French Infrastructure for Integrated Structural Biology FRISBI [ANR-10-INBS-05]. Figures 8, 12, 13 were prepared using *Pymol* (DeLano, 2002).

## Appendix A. Charge of atomic images

With the dimensionless parameter **x** = *D*^−1^**r**, and with *t* = 2*πx* = 2*π*|**x**|, the ‘charge’ of the image of the point atom (14) inside the sphere |**r**| ≤ *R* is calculated as

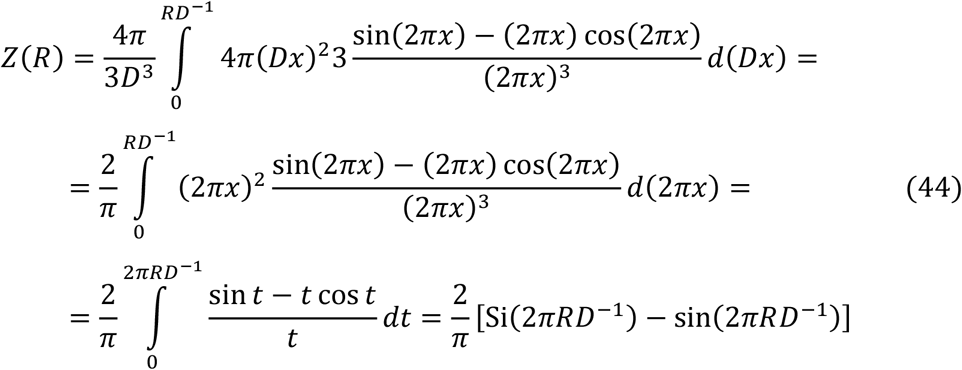

For *R* → ∞, the first term in the brackets, the sine integral, converges to 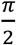, the desired answer. However, the second term of *Z*(*R*) oscillates around zero with the amplitude 1. The difference from the correct value is minimal when *RD*^−1^ is integer or half-integer.

## Appendix B. Test macromolecular models

The protein models used for the tests were constructed from the PDB entry 1zud already used previously (Urzhumtsev *et al*., 2014). After excluding all water molecules and alternative conformations, the model contained about 4500 atoms of four types: C, N, O, S. Atomic displacement factors *B_n_* of the atoms were taken as they are in the reported model; they varied from 7 to 97 Å^2^ with the mean value of 33 Å^2^. The model was placed in a virtual unit cell in space group P1 with the unit cell size *a* = 60 Å, *b* = 72 Å, *c* = 80 Å, *α* = *β* = *γ* = 90°; a single copy in the unit cell. This copy had weak contacts with its neighbors like in a real crystal. The 5-Gaussian approximation for X-ray scattering factors (Brown *et al*., 2006) were used.

For the same set of atomic coordinated, we prepared several more models, with a different degree of simplification of the atomic scattering factors:

- each *B_n_* value was replaced by the closest one from the grid 5, 10, 15, 20, 30, …, 100 Å^2^ (*B*-grid in the main text; 12 values in total);
- all *B_n_* values were kept as they are in the PDB model but all atoms were considered as carbons;
- two modifications above combined: all atoms were considered as carbons and their *B_n_* values replaced by the closest from the *B*-grid;
- all atoms are considered as carbons taken with *B_n_* value common for all atoms and varied from one test to another.

